# Nurr1 modulation mediates neuroprotective effects of statins

**DOI:** 10.1101/2021.09.15.460433

**Authors:** Sabine Willems, Whitney Kilu, Giuseppe Faudone, Jan Heering, Daniel Merk

## Abstract

The ligand-sensing transcription factor Nurr1 emerges as a promising therapeutic target for neurodegenerative pathologies but Nurr1 ligands for functional studies and therapeutic validation are lacking. Here we report pronounced Nurr1 modulation by statins for which clinically relevant neuroprotective effects have been demonstrated. Several statins directly affected Nurr1 activity in cellular and cell-free settings with low micromolar to sub-micromolar potencies. Simvastatin exhibited anti-inflammatory effects in astrocytes which were abrogated by Nurr1 knockdown. Differential gene expression analysis in native and Nurr1 silenced cells revealed strong proinflammatory effects of Nurr1 knockdown while simvastatin treatment induced several neuroprotective mechanisms via Nurr1, for example, in energy utilization and reduced apoptosis. These findings suggest Nurr1 involvement in the well-documented but mechanistically elusive neuroprotection by statins.

## Introduction

The ligand-activated transcription factor nuclear receptor related-1 (Nurr1, NR4A2)(Wang et al., 2003) is a constitutively active orphan nuclear receptor. It is considered as neuroprotective transcriptional regulator and ascribed high therapeutic potential in neurodegenerative diseases. Nurr1 is expressed in several neuronal cell populations with highest levels in dopaminergic neurons and thought to protect neurons against injury(Decressac et al., 2013). Neuronal Nurr1 knockout in mice produced a phenotype resembling Parkinson’s Disease (PD)(Decressac et al., 2013; Kim et al., 2015) and in the neurotoxin MPTP induced model of PD in rodents, Nurr1 was downregulated resulting in neuroinflammation and enhanced apoptosis of neuronal cells(Liu et al., 2017) while Nurr1 overexpression in the same model reduced motor impairment and spatial learning deficits(Liu et al., 2017). In experimental autoimmune encephalomyelitis (EAE), heterozygous Nurr1 knockout mice developed the disease faster than wild-type mice(Montarolo et al., 2015), while enhanced Nurr1 signaling reduced incidence and severity of EAE(Montarolo et al., 2014). Neuronal Nurr1 expression was also significantly downregulated in models of Alzheimer’s Disease (AD) in an age-dependent fashion(Moon et al., 2015, 2019) and the transcription factor was shown to protect against AD-related pathology including Aβ accumulation, neuronal loss and microglial activation in vivo(Moon et al., 2019). In line with these observations from rodent models, altered Nurr1 expression has been detected in human PD, AD and multiple sclerosis (MS) patients(Liu et al., 2017; Moon et al., 2015, 2019; Satoh et al., 2005) further highlighting the great neuroprotective potential of Nurr1(Jakaria et al., 2019) which may hence be a very attractive therapeutic target to treat neurodegenerative pathologies.

Despite this therapeutic promise, knowledge on Nurr1 function and ligands is still scarce. A few weak Nurr1 modulators have been discovered(Bruning et al., 2019; Kim et al., 2015; Munoz-Tello et al., 2020; Rajan et al., 2020; de Vera et al., 2019; Willems et al., 2020, 2021) such as the prostaglandins A1 and E1 as potential endogenous ligands(Rajan et al., 2020). The antimalarials amodiaquine (AQ) and chloroquine (CQ) have served as early Nurr1 agonist tools to evaluate Nurr1 activation in neurodegeneration(Kim et al., 2015, 2016; Moon et al., 2019). Therapeutic validation of Nurr1 in neurodegenerative pathologies and beyond, however, requires potent and selective Nurr1 modulators. Aiming to close this gap and expand the sparse collection of Nurr1 ligand scaffolds, we have screened a drug fragment library for Nurr1 modulation in a cellular setting resulting in the discovery of statins as potent Nurr1 modulators. Intrigued by this finding and reports on clinically relevant effects of this drug class in neurodegeneration(Carroll and Wyse, 2017; Chataway et al., 2014; Torrandell-Haro et al., 2020), we have evaluated the potential involvement of Nurr1 in the neuroprotective actions of statins. Differential gene expression experiments in native and Nurr1-silenced astrocytes demonstrated several Nurr1 mediated neuroprotective mechanisms of simvastatin indicating important contributions of Nurr1 modulation in the pharmacological effects of simvastatin and related drugs in neurodegeneration.

## Results

### Fragment screening reveals structurally diverse Nurr1 ligands

As rapid approach to discover Nurr1 ligands, we have screened a commercially available collection of 480 drug fragments (see Supplementary Figure 1 for details) for Nurr1 modulation in a cellular Gal4-Nurr1 hybrid reporter gene assay at a single concentration of 100 μM. Fragments affecting reporter activity ≥1.5-fold (Nurr1 activation) or ≤0.6-fold (Nurr1 repression) were considered for further evaluation (Figure 1 and Supplementary Figure 2A). Curation for toxicity and PAINs structures, and control experiments for non-specific effects on reporter activity (using Gal4-VP16(Budzyński et al., 2015; Sadowski et al., 1988)) resulted in a collection of seven Nurr1 ligand fragments with no privileged scaffold for further characterization. Four fragments promoted Nurr1 activity and three fragments acted as inverse Nurr1 agonists (Supplementary Figure 2). 3-(4-Fluorophenyl)indole emerged as most active Nurr1 activator fragment (EC_50_ 8.2 μM, 2.1-fold eff.). It is contained in the widely used cholesterol-lowering drug fluvastatin which was an even more potent Nurr1 agonist (EC_50_ 1.8 μM, 2.2-fold eff.). Following this remarkable finding, we tested all seven marketed statins (fluvastatin, lovastatin, simvastatin, pravastatin, atorvastatin, rosuvastatin, pitavastatin) for Nurr1 modulatory activity and observed Nurr1 agonism for all seven drugs except pravastatin (Figure 1B) but with differing potencies. Lovastatin and simvastatin demonstrated similar potencies as fluvastatin while rosuvastatin was less active. Atorvastatin weakly activated Nurr1 (1.4-fold eff.) with sub-micromolar potency (EC_50_ 0.78 μM) and pitavastatin evolved as the most potent Nurr1 agonist amongst statins (EC_50_ 0.12 μM, 1.7-fold eff.). Interestingly, statins share structural features with the known Nurr1 agonists AQ and CQ as illustrated by multiple alignment (Figure 1C) but exhibit remarkably higher potencies (up to 400-fold) on Nurr1 (Figure 1D).

**Figure 1.**
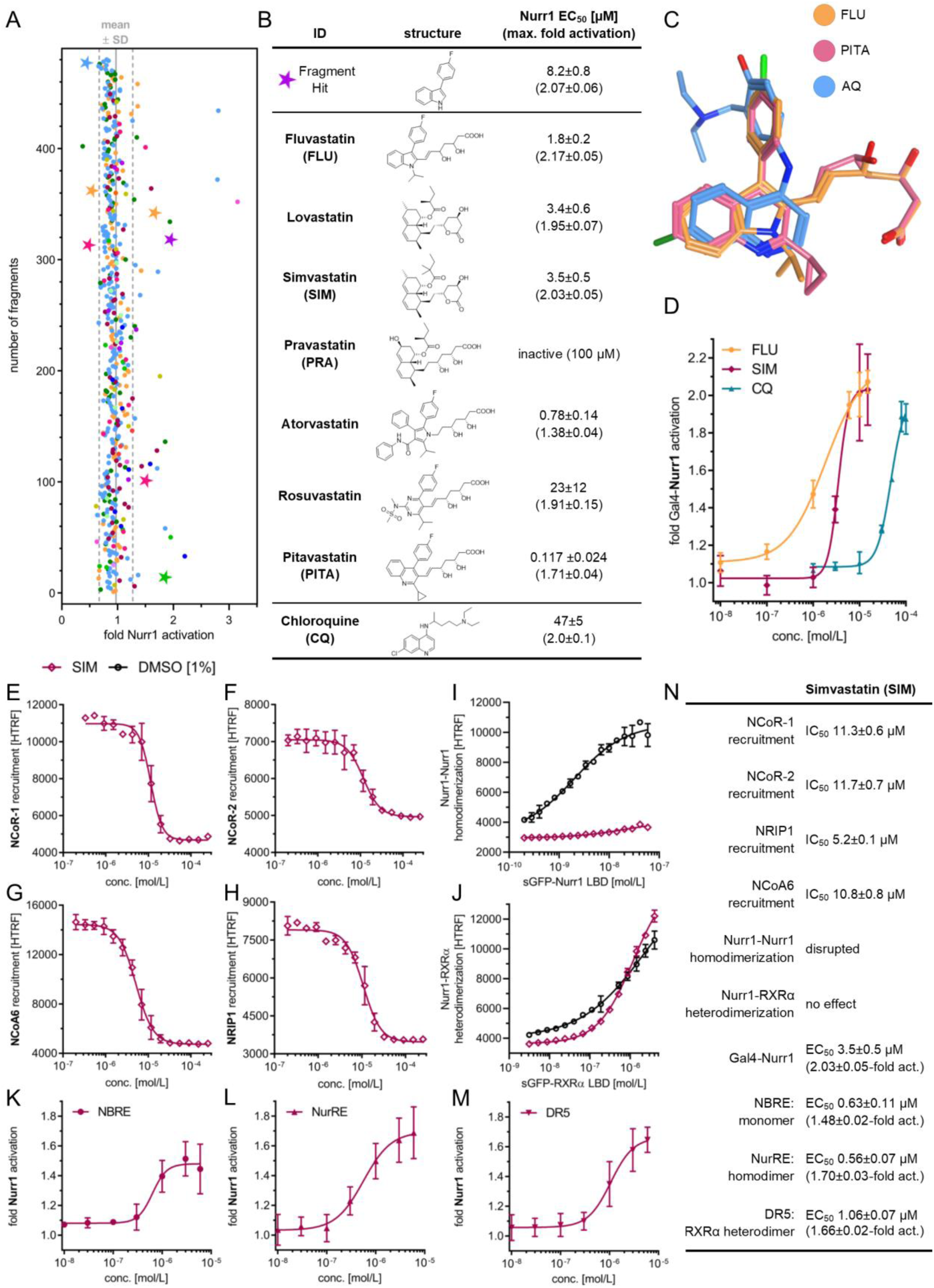
Discovery of statins as Nurr1 modulators and their profiling. (A) Primary fragment screening results. Nurr1 modulatory activity of the entire drug fragment library in a Gal4 reporter gene assay. Data are the mean reporter activity vs. 0.4% DMSO at 100 μM; n=2. Different colors represent different graph frameworks (see also Supplementary Figure 1). Compounds marked with a star relate to the fragment hits validated in control experiments on Gal4-VP16. Gray lines represent mean±SD of the entire screening. (B) Nurr1 modulatory activity of the fragment screening hit and of the statin class of drugs (vs. 0.1% DMSO) in a Gal4 hybrid Nurr1 reporter gene assay. Data are the mean±S.E.M.; n≥3. (C) Multiple alignment of fluvastatin (FLU), pitavastatin, and AQ reveals common structural features with overlap of the indole and quinoline scaffolds as well as the phenyl substituents. (D) Dose-response curves of simvastatin (SIM) and FLU determined in a Gal4-Nurr1 hybrid reporter gene assay. Chloroquine (CQ) for comparison. Data are the mean±S.E.M., n ≥ 3. (E-J) Effects of SIM on co-regulator interactions and dimerization of Nurr1 in HTRF assays. SIM displaced NCoR-1 (E), NCoR-2 (F), NCoA6 (G) and NRIP1 (H) from the Nurr1 LBD and SIM (30 μM) decreased homodimerization of Nurr1 (I) without affecting Nurr1-RXRα heterodimerization (J). Data are the mean±SD; N=3. (K-M) Profiling of SIM in human full-length Nurr1 reporter gene assays for the Nurr1 response elements NBRE (K, Nurr1 monomer), NurRE (L, Nurr1 homodimer), DR5 (M, Nurr1-RXRα heterodimer). Data are the mean±S.E.M., n≥3. (N) Summarized activities of Nurr1 modulator SIM in cell-free and cellular experiments.

### Statins modulate Nurr1 activity in cellular and cell-free settings

To further characterize the intriguing Nurr1 agonism of statins, we selected fluvastatin (highest Nurr1 agonist efficacy) and simvastatin (most widely used statin) as representative compounds. Additionally, simvastatin lacks a chromophore and was best suited for homogenous time-resolved fluorescence resonance energy transfer (HTRF) based assays. To obtain mechanistic insights in Nurr1 modulation by statins, we evaluated modulation of Nurr1 interactions with co-regulators by statins in cell-free HTRF based systems. We have previously discovered ligand-sensitive interaction of Nurr1 with nuclear receptor co-repressors (NCoR) 1 and 2, nuclear receptor interacting protein 1 (NRIP1) and nuclear receptor co-activator 6 (NCoA6)(Willems et al., 2020). Fluvastatin and simvastatin caused a concentration dependent displacement of all four co-regulators (Figure 1E-H, Supplementary Figure 3). Moreover, since Nurr1 can act as monomer, homodimer and RXR-heterodimer on different DNA response elements, its activity also depends on its dimerization state(Jiang et al., 2019; Willems et al., 2020). Fluvastatin and simvastatin did not alter heterodimerization of Nurr1 with RXRα but robustly inhibited Nurr1 homodimerization (Figure 1I,J, Supplementary Figure 3). The HTRF assays revealed higher potency of simvastatin compared to fluvastatin prompting us to perform further experiments with simvastatin. Next, we characterized the ability of simvastatin to modulate full-length human Nurr1 on the human monomer (NGFI-B response element, NBRE), homodimer (Nur-response element, NurRE), and heterodimer (direct repeat 5, DR5) response elements(Jiang et al., 2019). Simvastatin activated the full-length human Nurr1 on all three response elements with low micromolar to sub-micromolar potencies (Figure 1K-N). Despite disrupting Nurr1 homodimerization, simvastatin also activated the homodimer response element NurRE. As NurRE naturally also contains a Nurr1 monomer binding site(Maira et al., 1999; Willems et al., 2020), this finding is not surprising, however. A selectivity screen over lipid sensing nuclear receptors revealed no other activities of simvastatin and fluvastatin (Supplementary Figure 3G).

### Statins block the inflammatory response of astrocytes

To probe the relevance of Nurr1 modulation by statins in neuroinflammation, we studied the effects of simvastatin and fluvastatin on interleukin-6 (IL-6) release by Nurr1 expressing human astrocytes (T98G) in response to LPS treatment. Pravastatin, which does not activate Nurr1, was used as negative control. Simvastatin and fluvastatin markedly diminished LPS-induced IL-6 release while pravastatin had no effect suggesting Nurr1 involvement (Figure 2A). Silencing of Nurr1 by RNAi in T98G cells (Figure 2B) remarkably increased IL-6 production and abrogated the effect of simvastatin on IL-6 levels (Figure 2C) further supporting Nurr1 mediated activity of the statins.

**Figure 2.**
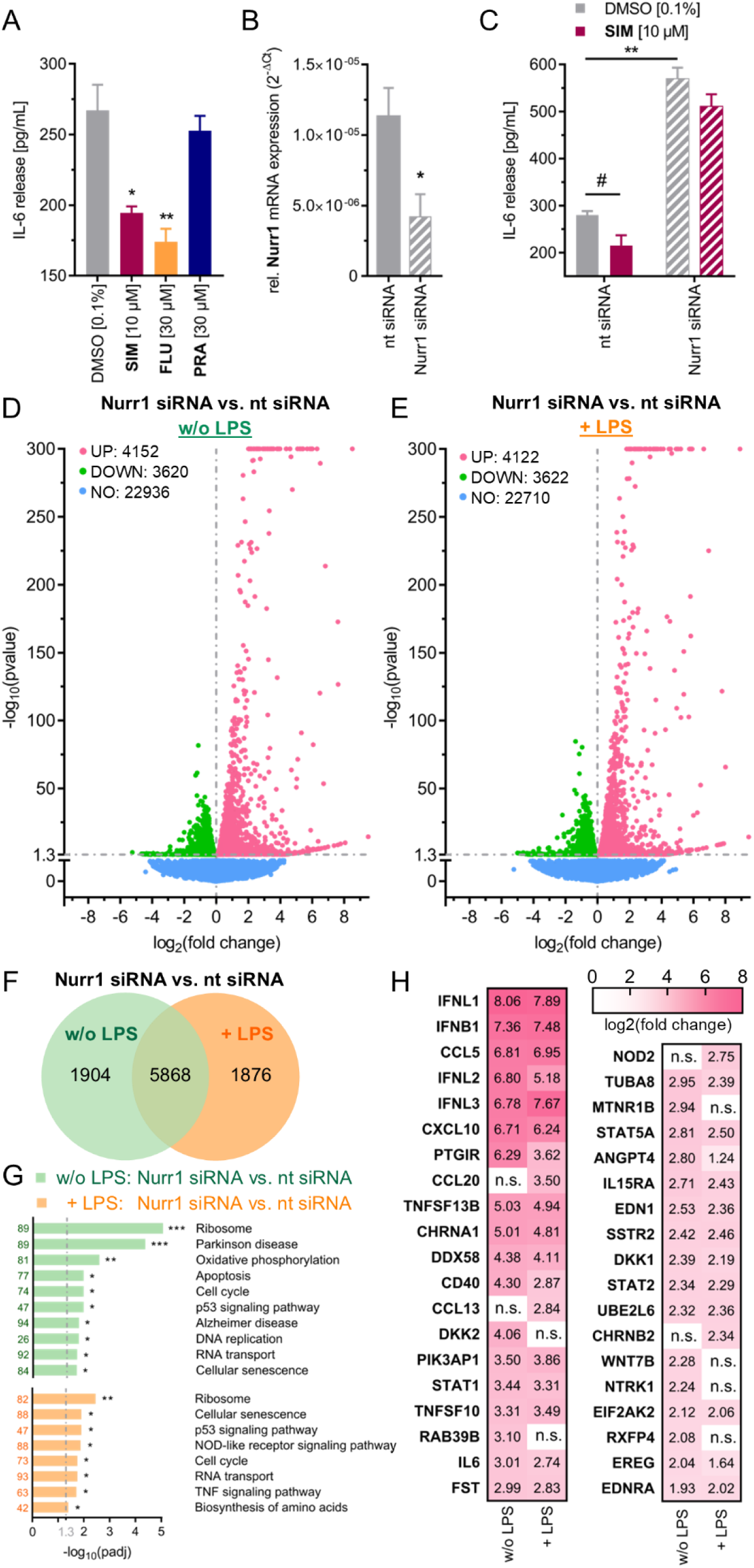
Nurr1 is involved in neuroinflammatory signaling. (A) The Nurr1 agonists SIM and FLU significantly countered interleukin-6 (IL-6) release from LPS-treated (1 μg/mL) T98G cells, whereas pravastatin (PRA), as negative control, did not affect IL-6 release. Data are the mean±S.E.M., n=4, ^#^ p < 0.1, * p < 0.05, ** p < 0.01 (t-test). (B) Nurr1 knockdown efficiency as determined by Nurr1 mRNA levels (qRT-PCR, 2^-ΔCt^ method with GAPDH as reference gene). Data are the mean±S.E.M., n=8, * p < 0.05 vs. non-targeting (nt) control siRNA (t-test). (C) LPS-treated T98G cells released considerable amounts of IL-6 which was further enhanced by siRNA-mediated Nurr1 knockdown suggesting reverse Nurr1 involvement in this inflammatory response. The Nurr1 agonist SIM ameliorated the inflammatory response of T98G cells in a Nurr1 dependent manner. Data are the mean±S.E.M., n=3, ^#^ p < 0.1, ** p < 0.01 (t-test). (D & E) Differential gene expression in T98G cells treated with nt or Nurr1 siRNA in absence (D) or presence (E) of LPS. Volcano plots show log_2_fold change in gene expression level (x axis) versus statistical significance level (-log_10_(p value); y axis). (F) Co-expression Venn diagram for differential gene expression in Nurr1 silenced cells versus nt siRNA for +/-LPS treated cells. (G) KEGG pathway enrichment analysis illustrates involvement of Nurr1 in signaling pathways related to neurodegenerative diseases and neuroinflammation. Bar plot shows statistical significance level (-log_10_(padj)) of regulated KEGG pathways, numbers refer to the count of differentially expressed genes related to the pathway. n=3, * p < 0.05, ** p < 0.01, *** p < 0.001. (H) Differentially expressed genes with log_2_fold change > |2| associated with neurodegenerative diseases (PD, AD, neurodegeneration) or neuroinflammatory signaling (TNF, NFκB, WNT, TGFβ, JAK-STAT, PI3K-Akt, apoptosis, neuroactive interaction) according to KEGG are listed with their respective log_2_fold change values for Nurr1 silencing compared to nt siRNA control in absence or presence of LPS. n.s. – not significant.

### Nurr1 knockdown alters neuroinflammatory signaling in vitro

The pronounced effect of Nurr1 knockdown on IL-6 levels aligned with the transcription factor’s important role in neuroprotection and -inflammation. To obtain insights in its neuroprotective mechanisms in astrocytes, we studied differential gene expression of T98G cells treated with Nurr1 siRNA or non-targeting (nt) siRNA in presence or absence of LPS (Figure 2D-H). siRNA mediated Nurr1 knockdown altered the expression of almost 8000 genes in both untreated and LPS-treated T98G cells but with pronounced differences of almost 2000 genes differentially affected by Nurr1 knockdown in presence or absence of LPS (Figure 2F). Silencing of Nurr1 strongly increased expression of multiple cytokines (interferons, C-C and C-X-C chemokines), cytokine receptors, TNF superfamily genes (e.g., CD40), and members of JAK-STAT signaling (STAT1, STAT2) even in absence of LPS stimulation (Figure 2H) indicating that diminished Nurr1 activity is sufficient to induce neuroinflammation. Still, LPS treatment caused upregulation of additional cytokines (CCL13, CCL20) in Nurr1 silenced cells. Pathway analysis of altered gene expression levels additionally revealed strong effects of Nurr1 silencing on genes involved in PD and AD, oxidative phosphorylation, apoptosis, and p53 signaling (Figure 2G).

### Simvastatin exhibits neuroprotective effects via Nurr1

As simvastatin had exhibited potential neuroprotective effects in LPS-treated astrocytes and significantly decreased IL-6 release, we evaluated its effect on gene expression in presence or absence of Nurr1. We treated astrocytes (T98G) with nt siRNA or Nurr1 siRNA and with DMSO or Simvastatin, stimulated the cells with LPS and determined differential gene expression (Figure 3). Compared to DMSO, simvastatin treatment had a pronounced effect on gene expression in both groups (nt and Nurr1 siRNA, Figure 3A,B). The expression levels of 1322 genes were affected by simvastatin treatment in both groups (Figure 3C) indicating that also other pathways were involved. 1389 genes were selectively affected by simvastatin (vs. DMSO) in nt siRNA treated cells but not altered upon simvastatin treatment of Nurr1 silenced cells (Figure 3B,C). Pathway analysis of nt siRNA vs. Nurr1 siRNA treated cells demonstrated Nurr1 mediated effects of simvastatin on genes related to PD, AD, apoptosis, p53 signaling, cell cycle and oxidative phosphorylation (Figure 3D). 109 genes revealed a strong response to simvastatin only in presence of Nurr1 with > |0.5| log2fold change (Table 1). Closer inspection of these Nurr1 mediated simvastatin effects revealed several neuroprotective effects including pronounced induction of hexokinase 3 (HK3), the E3-ubiquitin ligases RING finger protein RNF43 and RNF222, and notch4 as well as downregulation of gasdermin C (Figure 3E). HK3 is the rate-limiting enzyme of glucose utilization and its induction by simvastatin may importantly contribute to neuroprotective effects. Energy metabolism is critical for neuronal health and function, and altered glucose utilization in brain has been linked to neurodegenerative diseases, particularly to AD(Cisternas et al., 2016; Haenig et al., 2020; Winkler et al., 2015). Additionally, HK3 has been associated with cytoprotective effects against oxidative stress, increased ATP levels and enhanced mitochondrial biogenesis(Wyatt et al., 2010). RNF43 is considered as an anti-apoptotic regulator(Shinada et al., 2011), as a Wnt antagonist(Zhong et al., 2021) and to be involved in DNA repair(Lerksuthirat et al., 2020) suggesting its upregulation as another neuroprotective contribution. Moreover, despite incomplete understanding of notch in neurodegeneration(Ables et al., 2011), decreased notch signaling has been detected in AD(Alberi et al., 2013; Moehlmann et al., 2002) indicating a potential benefit of notch induction by simvastatin. Gasdermin C is a membrane pore-forming protein and a key mediator of inflammation and cell death(Broz et al., 2020; Feng et al., 2018; Rogers et al., 2019). Gasdermin pores permeabilize cell membranes and damage mitochondria to release cytochrome C leading to inflammasome activation and enhanced apoptosis(Broz et al., 2020; Feng et al., 2018; Rogers et al., 2019). The pronounced Nurr1 mediated downregulation of gasdermin C by simvastatin potentially emerges as a key neuroprotective effect preventing inflammation, apoptosis and neuronal cell death. In addition, marked Nurr1 mediated effects of simvastatin on neurotransmitter receptors and transporters as well as on inflammatory genes were evident from the comparison of nt siRNA and Nurr1 siRNA treated cells. Simvastatin downregulated metabotropic glutamate receptor 4 (GRM4), GABA receptor A3 (GABRA3) and the neuropeptide PEN receptor GPR83 while no neurotransmitter or neuropeptide receptor was induced. Simvastatin treatment also diminished expression of the GABA transporter SLC6A12 and several ion channels (KCNA7, KCNB1, TRPV2). Nurr1 mediated effects of simvastatin on genes involved in inflammation included induction of NFκB inhibitor alpha (NFKBIA) and intercellular adhesion molecule 1 (ICAM1), and downregulation of arachidonate 12 lipoxygenase (ALOX12) and IL-31 receptor (IL31RA). Cyclooxygenase 2 (COX-2, PTGS2) was upregulated suggesting that Nurr1 activation did not fully block LPS effects. Overall, differential gene expression analysis demonstrated distinguished Nurr1 mediated neuroprotective effects of simvastatin with anti-apoptotic, metabolic and anti-inflammatory contributions.

**Figure 3.**
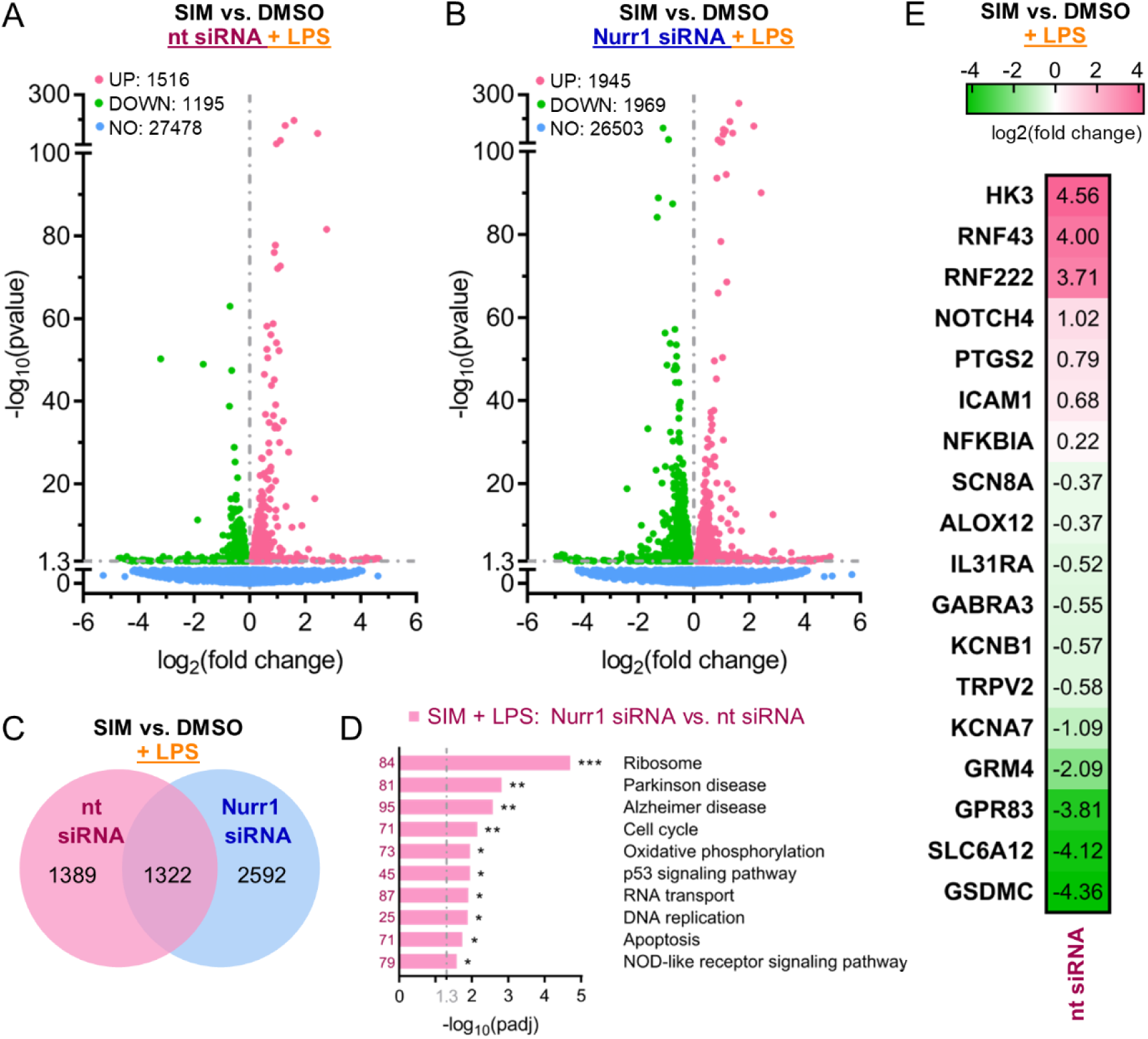
Simvastatin affected gene expression of LPS-treated human astrocytes (T98G) in a Nurr1 dependent manner. (A & B) Differentially expressed genes for SIM (10 μM) versus DMSO in LPS-stimulated T98G cells treated with nt siRNA (A) or Nurr1 siRNA (B). Volcano plots show log_2_fold change in gene expression level (x axis) versus statistical significance level (-log_10_(p value); y axis), n=3. (C) Co-expression Venn diagram shows effects of SIM vs. DMSO in LPS-stimulated T98G cells treated with nt siRNA (magenta) or Nurr1 siRNA (blue). (D) KEGG pathway enrichment analysis illustrates involvement of Nurr1 activation by SIM in signaling pathways related to neurodegeneration and neuroinflammation. Bar plot shows statistical significance level (-log_10_(padj)) of regulated KEGG pathways, numbers refer to the count of differentially expressed genes related to the pathway. n=3, * p < 0.05, ** p < 0.01, *** p < 0.001. (E) Genes regulated by SIM (vs. DMSO) in LPS stimulated nt siRNA treated cells whose expression was unaffected in Nurr1 siRNA treated cells. Only selected genes related to neuroprotection and neuroinflammation are shown, further regulated genes in Table 1. Heatmap shows log_2_fold change in gene expression.

**Table 1.**
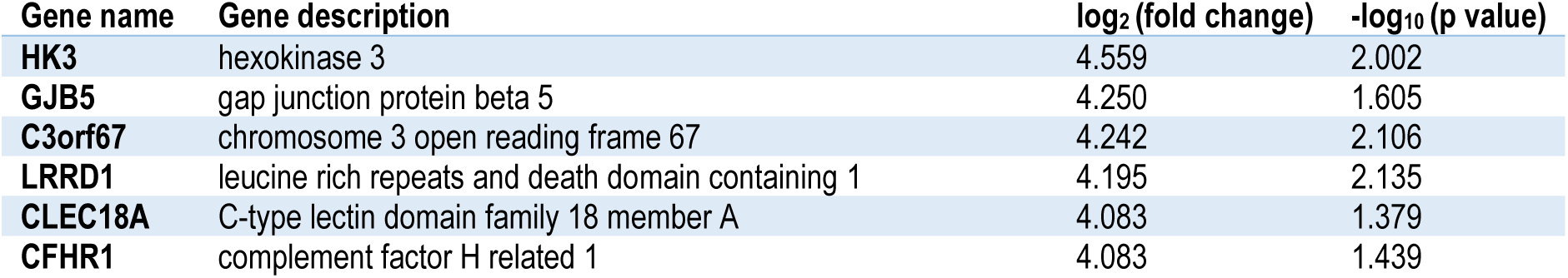
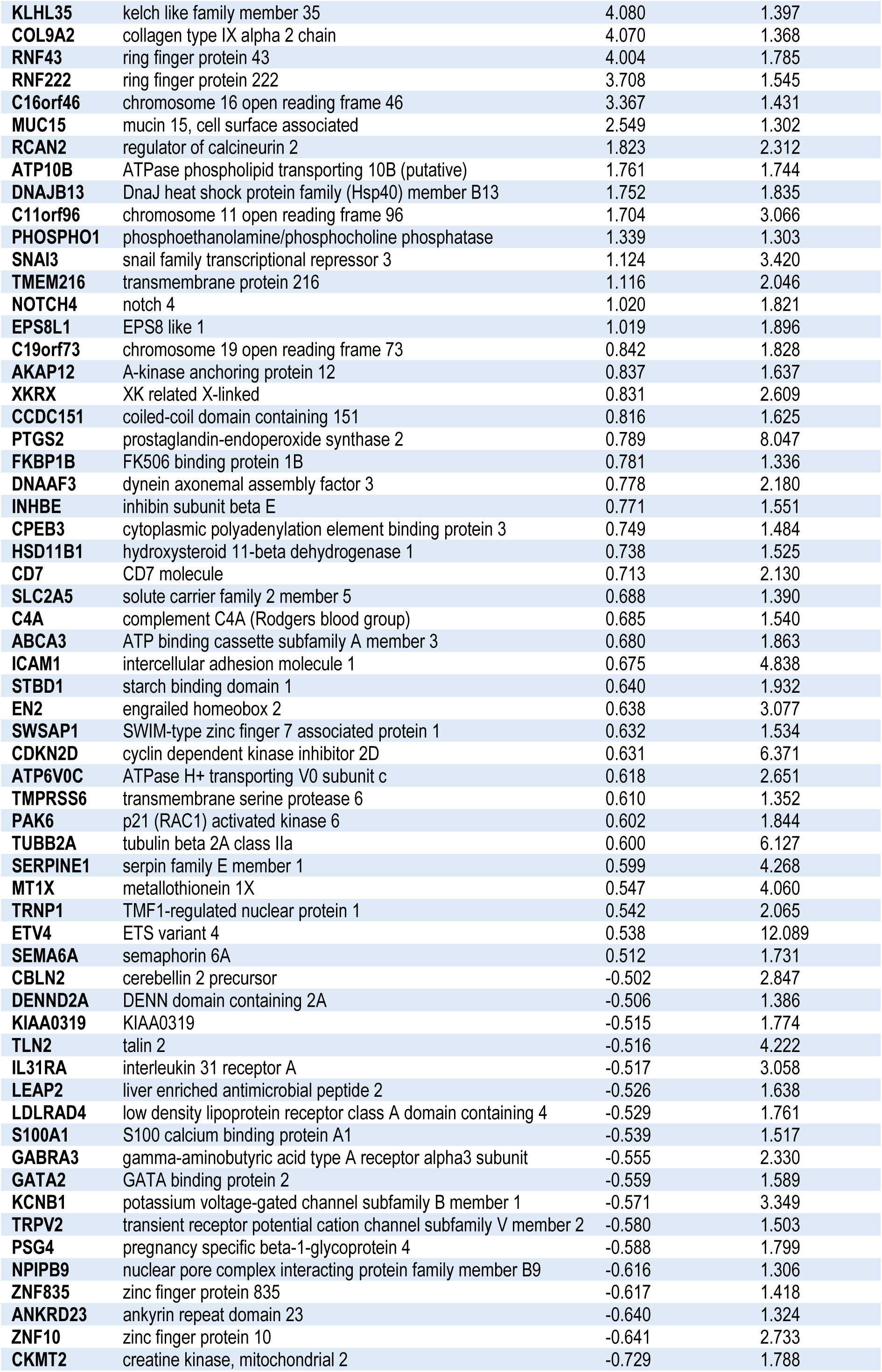
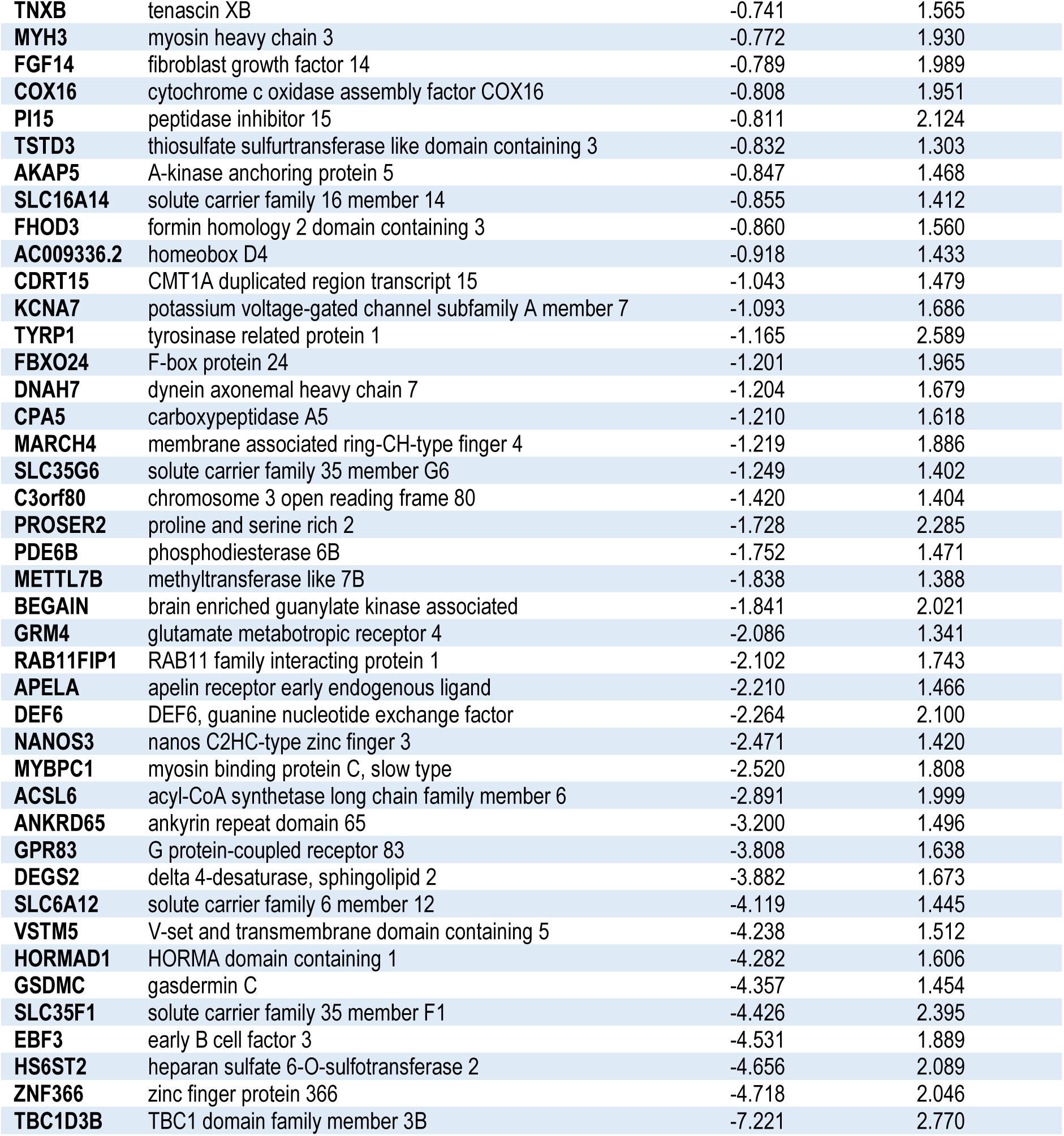
Effects of Simvastatin on differential gene expression in LPS-stimulated T98G cells. Only protein coding genes with log_2_fold change > |0.5| that were selectively altered by Simvastatin in nt siRNA treated cells but not altered in Nurr1 silenced cells are shown.

## Discussion

Several lines of evidence point to an important role and great therapeutic potential of the ligand sensing transcription factor Nurr1 in AD, PD and MS but pharmacological validation and exploitation of Nurr1 as therapeutic target is pending. In an attempt to rapidly expand the knowledge on Nurr1 ligand scaffolds, we have screened a drug fragment collection for Nurr1 modulation and discovered remarkable Nurr1 agonism of statins. Simvastatin activated the human Nurr1 on all its three human response elements with low micromolar to sub-micromolar potency and comprehensive mechanistic characterization revealed displacement of NCoR-1, NCoR-2, NRIP1 and NCoA6 from the Nurr1 LBD, and decreased Nurr1 homodimerization as key contributions to statin dependent Nurr1 activation.

The unprecedented molecular activity of the widely used drug class of statins on Nurr1 intriguingly aligns with several reports on neuroprotective effects of statins and suggests a potential involvement of Nurr1. Indeed, simvastatin counteracted inflammation in Nurr1 expressing astrocytes while this effect was lost in cells silenced for Nurr1 confirming relevance of Nurr1 activation by statins in neuronal cells. To capture neuronal effects of Nurr1 modulation by statins, we studied differential gene expression in Nurr1 expressing and silenced astrocytes upon simvastatin treatment. These experiments interestingly revealed strong pro-inflammatory effects of Nurr1 knockdown with strongly increased expression of multiple cytokines. Even more intriguing, our results demonstrate Nurr1 mediated neuroprotective effects of simvastatin including, for example, enhanced glucose utilization, altered notch signaling and several anti-apoptotic mechanisms that were not significantly affected in Nurr1 silenced cells. Of note, simvastatin (and other statins) crosses the blood-brain-barrier(Johnson-Anuna et al., 2005) supporting a potential clinical relevance of Nurr1 activation in the CNS by statins. Hence, our findings suggest that Nurr1 activation – together with other confirmed mechanisms(Ghosh et al., 2009; Huang et al., 2017; van der Most et al., 2009; Xu et al., 2013; Yan et al., 2020) – is involved in the neuroprotective effects of statins.

Protective and therapeutic effects of statin treatment in neurodegenerative diseases have been reported by several studies(Carroll and Wyse, 2017; Chataway et al., 2014; Torrandell-Haro et al., 2020). Particularly the use of simvastatin has been correlated with a suppression of proinflammatory molecules and microglial activation, inhibition of oxidative stress and attenuation of alpha-synuclein aggregation(Carroll and Wyse, 2017). Important clinical evidence for therapeutic potential of statins was described by Wahner et al. who found protective effects for all statins except pravastatin against PD(Wahner et al., 2008) which is particularly notable since pravastatin as only statin also failed to activate Nurr1. Observations on promising therapeutic potential in neurodegenerative diseases have evoked interventional clinical trials to reveal efficacy of simvastatin treatment in AD, PD and MS. While the PD-STAT(Carroll et al., 2019) trial could not confirm that simvastatin slows PD progression(Leigh, 2020), impressive results on efficacy of simvastatin in secondary progressive MS have been reported from the MS-STAT phase 2 trial(Chan et al., 2017; Chataway et al., 2014). Daily simvastatin treatment over two years significantly reduced brain atrophy compared to placebo and improved frontal lobe function and physical quality-of-life. The study concluded that its results support phase 3 testing but also noted that the mode-of-action for the observed neuroprotective effects of simvastatin in MS remains to be established. While HMG-CoA reductase inhibition and improved cholesterol balance undoubtedly contribute to neuroprotective statin effects, there are also cholesterol-independent activities the biochemical mechanisms of which remained elusive. Here, Nurr1 activation by statins evolves as a potentially critical mechanistic aspect in neuroprotective statin actions.

### Experimental Procedures

#### Chemicals and compounds

All compounds tested in this study were obtained from commercial vendors (Prestwick Chemical Libraries, Illkirich, France; TCI Chemicals Deutschland GmbH, Eschborn, Germany; Sigma Aldrich, St. Louis, MO, U.S.A.; Alfa Aesar, Ward Hill, MA, U.S.A.; abcr GmbH, Karlsruhe, Germany; Cayman Chemical, Ann Arbor, MI, U.S.A.; Fluorochem Ltd., Glossop, United Kingdom; Oxchem Corp., Wood Dale, IL, U.S.A.).

#### Hybrid reporter gene assays

##### Plasmids

The Gal4-fusion receptor plasmids pFA-CMV-hNURR1-LBD, pFA-CMV-hPPARα-LBD, pFA-CMV-hPPARγ-LBD, pFA-CMV-hPPARδ-LBD, pFA-CMV-hRARα-LBD and pFA-CMV-hRXRα-LBD coding for the hinge region and LBD of the canonical isoform of the respective human nuclear receptor have been reported previously(Willems et al., 2020). The Gal4-VP16(Sadowski et al., 1988) fusion protein expressed from plasmid pECE-SV40-Gal4-VP16(Budzyński et al., 2015) (Addgene, entry 71728, Watertown, MA, USA) served as ligand-independent transcriptional inducer for control experiments. pFR-Luc (Stratagene, La Jolla, CA, USA) was used as reporter plasmid and pRL-SV40 (Promega, Madison, WI, USA) for normalization of transfection efficiency and test compound toxicity. *Assay procedure*: HEK293T cells were grown in DMEM high glucose, supplemented with 10% fetal calf serum (FCS), sodium pyruvate (1 mM), penicillin (100 U/mL) and streptomycin (100 μg/mL) at 37 °C and 5% CO_2_. The day before transfection, HEK293T cells were seeded in 96-well plates (3 × 10^4^ cells/well). Medium was changed to Opti-MEM without supplements right before transfection. Transient transfection was performed using Lipofectamine LTX reagent (Invitrogen, Carlsbad, CA, USA) according to the manufacturer’s protocol with pFR-Luc (Stratagene), pRL-SV40 (Promega) and the corresponding Gal4-fusion nuclear receptor plasmid pFA-CMV-hNR-LBD. 5 h after transfection, medium was changed to Opti-MEM supplemented with penicillin (100 U/mL) and streptomycin (100 μg/mL), now additionally containing 0.1% DMSO and the respective test compound or 0.1% DMSO alone as untreated control. For the primary screen, each concentration was tested as single point measurements and each experiment was performed independently two times with 0.4% DMSO, respectively. For full dose-response characterization, each concentration was tested in duplicates and each experiment was performed independently at least three times. The Gal4-VP16 control experiment was carried out in duplicates as well, with at least four independent repeats. Following overnight (12-14 h) incubation with the test compounds, cells were assayed for luciferase activity using Dual-Glo™ Luciferase Assay System (Promega) according to the manufacturer’s protocol. Luminescence was measured with a Spark 10M luminometer (Tecan Group AG, Männedorf, Switzerland). Normalization of transfection efficiency and cell growth was done by division of firefly luciferase data by renilla luciferase data and multiplying the value by 1000 resulting in relative light units (RLU). Fold activation was obtained by dividing the mean RLU of a test compound at a respective concentration by the mean RLU of untreated control. Max. relative activation refers to fold reporter activation of a test compound divided by the fold activation of the respective reference agonist (PPARα: GW7647; PPARγ: rosiglitazone; PPARδ: L165,041; RXRα: bexarotene; RARα: tretinoin; all at a concentration of 1 μM; Nurr1: amodiaquine (100 μM)). All hybrid assays were validated with the above mentioned reference agonists which yielded EC_50_ values in agreement with the literature. For dose-response curve fitting and calculation of EC_50_/IC_50_ values, the equations “[Agonist]/[Inhibitor] vs. response – Variable slope (four parameters)” were performed with mean fold activations ± S.E.M. using GraphPad Prism (version 7.00, GraphPad Software, La Jolla, CA, USA).

#### Reporter gene assays involving full-length human Nurr1

##### Plasmids

The reporter plasmids pFR-Luc-NBRE(Willems et al., 2020), pFR-Luc-NurRE(Willems et al., 2020) and pFR-Luc-DR5(Willems et al., 2020) each containing one copy of the respective human Nurr1 response element NBRE Nl3 (TGATATCGAAAACA<u>AAAGGTCA</u>), NurRE (from POMC; <u>TGATATTTACCTCCAAATGCCA</u>) or DR5 (TGATA<u>GGTTCACCGAAAGGTCA</u>), were described previously. The full length human nuclear receptor Nurr1 (pcDNA3.1-hNurr1-NE; Addgene, entry 102363) and, for DR5, RXRα (pSG5-hRXR) were overexpressed. pFL-SV40 (Promega) was used for normalization of transfection efficacy and evaluation of compound toxicity. *Assay procedure*: HEK293T cells were grown in DMEM high glucose, supplemented with 10% FCS, sodium pyruvate (1 mM), penicillin (100 U/mL) and streptomycin (100 μg/mL) at 37 °C and 5% CO_2_. The day before transfection, HEK293T cells were seeded in 96-well plates (3 × 10^4^ cells/well). Medium was changed to Opti-MEM without supplements right before transfection. Transient transfection was performed using Lipofectamine LTX reagent (Invitrogen) according to the manufacturer’s protocol with pFR-Luc-NBRE(Willems et al., 2020), pFR-Luc-NurRE(Willems et al., 2020) or pFR-Luc-DR5(Willems et al., 2020), pRL-SV40 (Promega), the human full length receptor plasmid pcDNA3.1-hNurr1-NE, and, for DR5, also pSG5-hRXR. 5 h after transfection, medium was changed to Opti-MEM supplemented with penicillin (100 U/mL) and streptomycin (100 μg/mL), now additionally containing 0.1% DMSO and the respective test compound or 0.1% DMSO alone as untreated control. For full dose-response characterization, each concentration was tested in duplicates and each experiment was performed independently at least three times. Following overnight (12-14 h) incubation with the test compounds, cells were assayed for luciferase activity using Dual-Glo™ Luciferase Assay System (Promega) according to the manufacturer’s protocol. Luminescence was measured with a Spark 10M luminometer (Tecan Group AG). Normalization of transfection efficiency and cell growth was done by division of firefly luciferase data by renilla luciferase data and multiplying the value by 1000 resulting in relative light units (RLU). Fold activation was obtained by dividing the mean RLU of a test compound at a respective concentration by the mean RLU of untreated control. The full length Nurr1 reporter gene assays were validated with amodiaquine and chloroquine as reference agonists. For dose-response curve fitting and calculation of EC_50_ values, the equation “[Agonist] vs. response – Variable slope (four parameters)” was performed with mean fold activations ± S.E.M. using GraphPad Prism (version 7.00, GraphPad Software).

#### Nurr1 co-regulator recruitment assays

Recruitment of co-regulator peptides to the Nurr1-LBD was studied in a homogeneous time-resolved fluorescence resonance energy transfer (HT-FRET) assay system. Terbium cryptate as streptavidin conjugate (Tb-SA; Cisbio Bioassays, Codolet, France) was used as FRET donor for stable coupling to biotinylated recombinant Nurr1-LBD protein(Willems et al., 2020) which has been reported previously. Four co-regulator peptides fused to fluorescein as FRET acceptor were purchased from ThermoFisher Scientific (Life Technologies GmbH, Darmstadt, Germany). Assay solutions were prepared in HTRF assay buffer (25 mM HEPES pH 7.5, 150 mM KF, 5% (w/v) glycerol, 5 mM DTT) supplemented with 0.1% (w/v) CHAPS and contained recombinant biotinylated Nurr1-LBD (3 nM), Tb-SA (3 nM) and the respective fluorescein-labeled co-regulator peptide (100 nM) as well as 1% DMSO with varying concentrations of the test compounds simvastatin or fluvastatin, or DMSO alone as negative control. All HTRF experiments were carried out in 384 well format using white flat bottom polystyrol microtiter plates (Greiner Bio-One, Frickenhausen, Germany). All samples were set up in triplicates. After 2 h incubation at RT, fluorescence intensities (FI) after excitation at 340 nm were recorded at 520 nm for fluorescein acceptor fluorescence and 620 nm for Tb-SA donor fluorescence on a SPARK plate reader (Tecan Group AG). FI520nm was divided by FI620nm and multiplied with 10,000 to give a dimensionless HTRF signal. The co-regulator peptides in this experiment were the following: nuclear receptor co-repressor (NCoR) 1, fluorescein-RTHRLITLADHICQIITQDFARN-OH; NCoR-2, fluorescein-HASTNMGLEAIIRKALMGKYDQW-OH; nuclear receptor co-activator 6 (NCoA6) fluorescein-VTLTSPLLVNLLQSDISAG-OH; nuclear receptor interacting protein 1 (NRIP1), fluorescein-SHQKVTLLQLLLGHKNEEN-OH. For dose-response curve fitting and calculation of IC_50_ values, the equation “[Inhibitor] vs. response – Variable slope (four parameters)” was performed with mean fold activations ± SD using GraphPad Prism (version 7.00, GraphPad Software).

#### Nurr1 dimerization assays

Modulation of Nurr1 LBD homodimerization and heterodimerization with RXRα LBD were studied in HT-FRET assay setups using biotinylated recombinant Nurr1-LBD(Willems et al., 2020) and GFP-Nurr1 LBD(Willems et al., 2020) or GFP-RXRα LBD(Willems et al., 2020), respectively. Assay solutions were prepared in HTRF assay buffer supplemented with 0.1% (w/v) CHAPS as well as 1% DMSO with test compounds at 30 μM or DMSO alone as negative control. The biotinylated Nurr1 LBD (0.375 nM) and Tb-SA (0.75 nM) served as FRET donor complex which was kept constant while the GFP-coupled protein as FRET acceptor was varied in concentration. Since affinity of both Nurr1 dimer formations differ, titration of GFP-Nurr1 LBD started at 500 nM and, for GFP-RXRα LBD, at 4 μM, respectively. Accordingly, free GFP was added to keep the total GFP content stable throughout the entire series in order to suppress artefacts from changes in degree of diffusion enhanced FRET. All samples were set up in triplicates and equilibrated at RT for 2 h before FI520 and FI620 were recorded after excitation at 340 nm, and the HTRF signal was calculated as described above.

#### Nurr1 knockdown in T98G cells

T98G cells (ATCC® CRL1690™) were grown in DMEM high glucose, supplemented with 10% FCS, sodium pyruvate (1 mM), penicillin (100 U/mL), and streptomycin (100 μg/mL) at 37°C and 5% CO_2_. 24 h before transfection, T98G cells were seeded in 12-well plates (1 × 10^5^ cells/well). The medium was changed to reduced serum medium containing DMEM high glucose supplemented with 1% charcoal-stripped FCS, sodium pyruvate (1 mM), penicillin (100 U/mL), and streptomycin (100 μg/mL) right before transfection. Knockdown was mediated by transient transfection using the RNAiMAX reagent (Invitrogen) according to the manufacturer’s protocol with 30 nM of Nurr1 targeting esiRNA (Cat# EHU008731) or non-targeting control siRNA (Cat# SIC001, both Sigma Aldrich). 24 h after transfection, the cells were rather harvested and directly used for RNA extraction or used for further experiments. 2 μg of total RNA were extracted from T98G cells by the E.Z.N.A. Total RNA Kit I (R6834-02, Omega Bio-Tek, Inc., Norcross, GA). RNA was reverse-transcribed into cDNA using the High-Capacity RNA-to-cDNA Kit (4387406, Thermo Fischer Scientific Inc., Waltham, MA, USA) according to the manufacturer’s protocol. Nurr1 knockdown efficiency was evaluated by quantitative real-time PCR (qRT-PCR) analysis with a StepOnePlus System (Life Technologies, Carlsbad, CA) using Power SYBR Green (Life Technologies; 12.5 μL/well). Each sample was set up in duplicates and repeated in eight independent experiments. The expression was quantified by the comparative 2^-ΔCt^ method and glyceraldehyde 3-phosphate dehydrogenase (GAPDH) served as the reference gene. Primer sequences for the human Nurr1 gene were obtained from OriGene (OriGene Technologies Inc., Rockville, MD, USA). The following PCR primers were used: hGAPDH: 5′-ATA TGA TTC CAC CCA TGG CA (fw), 5′-GAT GAT GAC CCT TTT GGC TC (rev), hNurr1: 5’-AAA CTG CCC AGT GGA CAA GCG T (fw), 5′-GCT CTT CGG TTT CGA GGG CAA A (rev).

#### Quantification of IL-6 Release in T98G cells

T98G (ATCC® CRL1690™) cells were grown in 12-well plates (1 × 10^5^ cells/well) for knockdown experiments or 24-well plates (5 × 10^4^ cells/well) in DMEM high glucose, supplemented with 10% FCS, sodium pyruvate (1 mM), penicillin (100 U/mL), and streptomycin (100 μg/mL) at 37 °C and 5% CO_2_ for 24 h. Before incubation with LPS and test compounds, the medium was changed to DMEM supplemented with 1% charcoal-stripped FCS, sodium pyruvate (1 mM), penicillin (100 U/mL), and streptomycin (100 μg/mL) for 12 h, or Nurr1 knockdown was performed by transient transfection for 24 h as outlined above. The cells were then treated with LPS (1 μg/mL) to induce inflammation and simultaneously incubated with simvastatin (10 μM), fluvastatin (30 μM) or pravastatin (30 μM), and 0.1% DMSO, or 0.1 % DMSO alone as untreated control. Each sample was repeated independently at least three times. Following overnight (12 h for knockdown or 24 h) incubation, 100 μL of the respective supernatants were collected and assayed for IL-6 using the Human IL-6 ELISA Kit (Cat# KHC0061, Thermo Fisher Scientific, Inc.) according to the manufacturer’s protocol. Absorbance at 450 nm was measured with a Spark 10 M luminometer (Tecan Group AG).

#### Differential gene expression analysis

##### Sample preparation

T98G cells (ATCC® CRL1690™) were cultured in DMEM, high glucose supplemented with 10% fetal calf serum (FCS), sodium pyruvate (1 mM), penicillin (100 U/mL), and streptomycin (100 μg/mL) at 37 °C and 5% CO_2_ and seeded in 12-well plates (1 × 10^5^ cells/well) for gene expression analysis. 24 h after seeding, medium was changed to reduced serum medium containing DMEM high glucose supplemented with 1% charcoal-stripped FCS, sodium pyruvate (1 mM), penicillin (100 U/mL), and streptomycin (100 μg/mL) right before transfection. Knockdown was mediated by transient transfection using the RNAiMAX reagent (Invitrogen) according to the manufacturer’s protocol with 30 nM of Nurr1 targeting esiRNA (Cat# EHU008731) or non-targeting control siRNA (Cat# SIC001, both Sigma Aldrich). After further 24 h, medium was changed again to reduced serum medium supplemented as described above now additionally containing 0.1% DMSO and the test compound simvastatin (10 μM) or 0.1% DMSO alone as control. Additionally, LPS (1 μg/mL) was added simultaneously to induce inflammation in one treatment arm. Each condition was set up in three independent biological repeats (n=3). After 12 h incubation, cells were harvested, washed twice with cold phosphate buffered saline (PBS) and then directly used for RNA extraction by the E.Z.N.A.® Total RNA Kit I (R6834-02, Omega Bio-Tek Inc., Norcross, GA, USA). *mRNA sequencing*. A total amount of 1 μg RNA per sample was used as input material for the RNA sample preparations. Sequencing libraries were generated using NEBNext® Ultra™ RNA Library Prep Kit for Illumina® (New England Biolabs (NEB), Ipswich, MA, U.S.A.) following manufacturer’s recommendations and index codes were added to attribute sequences to each sample. Briefly, mRNA was purified from total RNA using poly-T oligo-attached magnetic beads. Fragmentation was carried out using divalent cations under elevated temperature in NEBNext First Strand Synthesis Reaction Buffer (5X). First strand cDNA was synthesized using random hexamer primer and M-MuLV Reverse Transcriptase (RNase H-). Second strand cDNA synthesis was subsequently performed using DNA Polymerase I and RNase H. Remaining overhangs were converted into blunt ends via exonuclease/polymerase activities. After adenylation of 3’ ends of DNA fragments, NEBNext Adaptor with hairpin loop structure were ligated to prepare for hybridization. In order to select cDNA fragments of preferentially 150∼200 bp in length, the library fragments were purified with AMPure XP system (Beckman Coulter, Beverly, USA). Then 3 μL USER Enzyme (NEB, USA) was used with size-selected, adaptor ligated cDNA at 37 °C for 15 min followed by 5 min at 95°C before PCR. Then PCR was performed with Phusion High-Fidelity DNA polymerase, Universal PCR primers and Index (X) Primer. At last, PCR products were purified (AMPure XP system) and library quality was assessed on the Agilent Bioanalyzer 2100 system. The clustering of the index-coded samples was performed on a cBot Cluster Generation System using PE Cluster Kit cBot-HS (Illumina) according to the manufacturer’s instructions. After cluster generation, the library preparations were sequenced on an Illumina NovaSeq 6000 platform and paired-end reads were generated. *Data analysis*. Raw data (raw reads) of FASTQ format were firstly processed through fastp(Chen et al., 2018). In this step, clean data (clean reads) were obtained by removing reads containing adapter and poly-N sequences and reads with low quality from raw data. At the same time, Q20, Q30 and GC content of the clean data were calculated. All the downstream analyses were based on the clean data with high quality. Downstream analysis was performed using a combination of programs including STAR(Dobin et al., 2013), HTseq(Anders et al., 2015), Cufflink(Trapnell et al., 2012) and wrapped scripts. Alignments were parsed using TopHat program(Trapnell et al., 2012) and differential expressions were determined through DESeq2(Love et al., 2014). KEGG enrichment analysis was implemented by the ClusterProfiler. Reference genome and gene model annotation files were downloaded from genome website browser (NCBI/UCSC/Ensembl) directly. Indexes of the reference genome were built using STAR and paired-end clean reads were aligned to the reference genome using STAR (v2.5). STAR used the method of Maximal Mappable Prefix (MMP) which can generate a precise mapping result for junction reads. HTSeq v0.6.1 was used to count the read numbers mapped of each gene. Reads per kilobase of exon model per million mapped reads (RPKM, considering the effect of sequencing depth and gene length for the reads count at the same time) of each gene was calculated based on the length of the gene and reads count mapped to this gene. Differential expression analysis between two conditions/groups (three biological repeats per condition) was performed using the DESeq2 R package (2_1.6.3). DESeq2 provides statistical routines for determining differential expression in digital gene expression data using a model based on the negative binomial distribution. The resulting p-values were adjusted using the Benjamini and Hochberg’s approach for controlling the False Discovery Rate (FDR). Genes with an adjusted P-value <0.05 found by DESeq2 were assigned as differentially expressed. Venn diagrams were prepared using the function vennDiagram in R based on the gene list for different group. We used clusterProfiler R package(Yu et al., 2012) to test the statistical enrichment of differential expression of genes in KEGG pathways using the KEGG database resource (http://www.genome.jp/kegg/)(Kanehisa et al., 2017). Correlations between individual samples were determined using the cor.test function in R with options set alternative =“greater” and method =“Spearman”. To identify the correlation between differences, different samples were clustered by expression level RPKM using hierarchical clustering distance method with the function heatmap, SOM (Self-organization mapping) and kmeans using silhouette coefficient to adapt the optimal classification with default parameter in R.

## Data and code availability

Differential gene expression data are available at ArrayExpress with the accession code E-MTAB-10624.

## Acknowledgements

D.M. is grateful for support by the Aventis Foundation. This work was supported by the research funding program LOEWE of the State of Hessen, Research Center for Translational Medicine and Pharmacology TMP.

## Abbreviations

AD: Alzheimer’s Disease
DR5: direct repeats spaced by 5 nucleotides
AQ: amodiaquine
EAE: experimental autoimmune encephalomyelitis
GFP: green fluorescent protein
HBA: number of hydrogen-bond acceptor
HBD: number of hydrogen-bond donor
HTRF: homogenous time-resolved fluorescence resonance energy transfer
KEGG: Kyoto encyclopedia of genes and genomes
LBD: ligand binding domain
MPTP: 1-Methyl-4-phenyl-1,2,3,6-tetrahydropyridin
MS: multiple sclerosis
NBRE: NGFI-B response element
NCoA6: nuclear receptor co-activator 6
NCoR-1: nuclear receptor co-repressor 1
NCoR-2: nuclear receptor co-repressor 2
NR: nuclear receptor
NRIP1: nuclear receptor interacting protein 1
NSAIDs: non-steroidal anti-inflammatory drugs
Nurr1: nuclear receptor related 1
NurRE: Nur-response element
PAINS: pan assay interference compounds
PD: Parkinson’s Disease
POMC: pro-opiomelanocortin
PPAR: peroxisome proliferator-activated receptor
qRT-PCR: quantitative real-time polymerase chain reaction
RAR: retinoic acid receptor
RT: room temperature
RXR: retinoid X receptor
SD: standard deviation
S.E.M.: standard error of the mean
TPSA: topological polar surface area
TR-FRET: time-resolved fluorescence resonance energy transfer.

## Supplementary Information

Figure S1. Characteristics of the fragment library and primary screening results.

Figure S2. Follow up of the primary drug fragment screen for Nurr1 modulation.

Figure S3. Effects of fluvastatin (FLU) on co-regulator interactions and dimerization of Nurr1.

Supplementary Methods: Computational Methods.

