## Supplementary Information for "Nurr1 modulation mediates neuroprotective effects of statins"

#### Table of contents

Supplementary Figures ........................................ 2

Supplementary Methods ........................................ 6

### Supplementary Figures

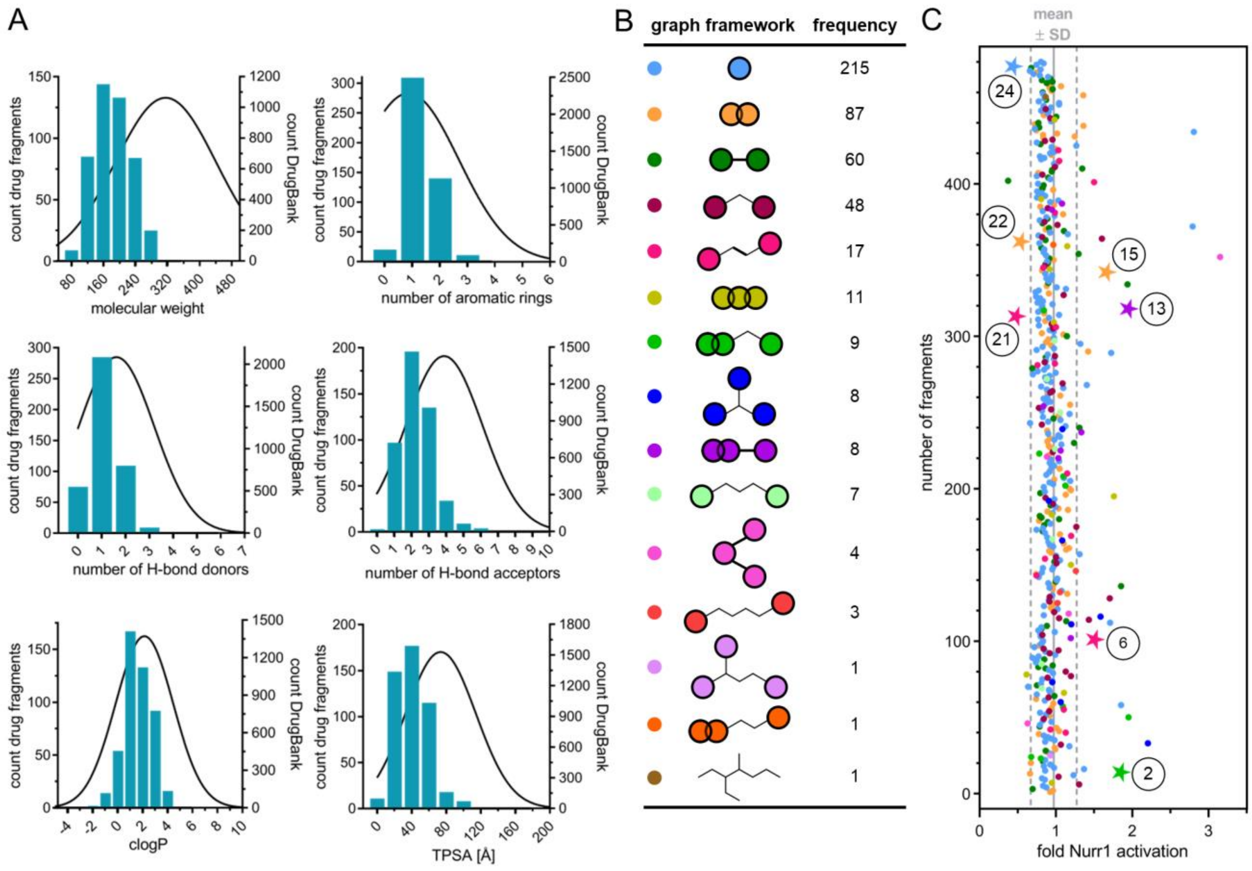


**Supplementary Figure 1.** (A) Feature distributions of the drug fragment library (N=480) in comparison with the DrugBank database (N=7946). Bars are plotted on the left y-axis for Prestwick drug fragments. Gaussian distributions of DrugBank compounds (MW ≤ 500) are plotted on the right y-axis. (B) Representations of the different graph frameworks contained in the drug fragment library and their frequencies. (C) Nurr1 modulatory activity of the drug fragment library in a Gal4 hybrid Nurr1 reporter gene assay. Results from primary screen are the mean reporter activity vs. 0.4% DMSO; n=2. Fragments affecting reporter activity ≥1.5-fold (Nurr1 activation, **1-20**) or ≤0.6-fold (Nurr1 repression, **21-24**) were considered for further evaluation as primary screening hits. Labeled compounds marked with a star relate to the fragment hits validated in control experiments on Gal4-VP16. Different colors refer to different graph frameworks (from B). Gray lines represent mean±SD of the entire screening.

A

| fragment hit | structure | primary screen (reporter act. at 100 µM) | Nurr1 modulation |
|---|---|---|---|
| **1** | 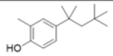 | 4.16±5.23 | toxic in primary screen |
| **2** | 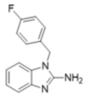 | 1.85±0.04 | $EC_{50}$ 7.2±1.1 (1.68±0.07 max. act.) |
| **3** | 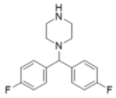 | 2.21±0.83 | inactive (50µM) toxic (≥ 100 µM) |
| **4** | 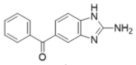 | 1.95±0.33 | inactive (50 µM) toxic (≥ 100 µM) |
| **5** | 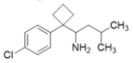 | 1.85±0.31 | inactive (30 µM) toxic (≥ 50 µM) |
| **6** | 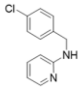 | 1.51±0.36 | $EC_{50}$ 16±1 (1.50±0.02 max. act.) |
| **7** | 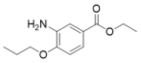 | 1.71±0.07 | inactive (50 µM) toxic (≥ 100 µM) |
| **8** | 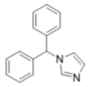 | 1.59±0.29 | inactive (30 µM) toxic (≥ 50 µM) |
| **9** | 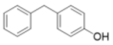 | 1.71±0.02 | toxic in primary screen |
| **10** | 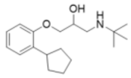 | 1.86±0.43 | inactive (30 µM) toxic (≥ 100 µM) |
| **11** | 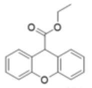 | 1.76±0.10 | inactive(50 µM) toxic (≥ 100 µM) |
| **12** | 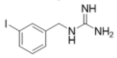 | 1.73±0.35 | toxic in primary screen |
| **13** | 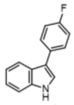 | 1.95±1.44 | $EC_{50}$ 8.2±0.8 (2.07±0.06 max. act.) |
| **14** | 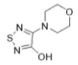 | 1.94±0.24 | inactive (10 µM) toxic (≥ 30 µM) |
| **15** | 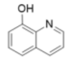 | 1.66±0.33 | $EC_{50}$ 22±2 (1.62±0.02 max. act.) |
| **16** | 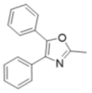 | 3.15±0.79 | inactive (10 µM) toxic (≥ 30 µM) |
| **17** | 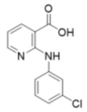 | 1.60±0.23 | inactive(10 µM) toxic (≥ 30 µM) |
| **18** | 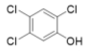 | 2.79±0.00 | toxic in primary screen |
| **19** | 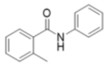 | 1.50±0.05 | inactive (100 µM) |
| **20** | 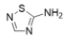 | 2.81±0.32 | toxic in primary screen |
| **21** | 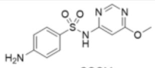 | 0.48±0.05 | $IC_{50}$ 35±3 (0.48±0.03 rem. act.) |
| **22** | 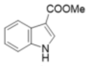 | 0.54±0.06 | $IC_{50}$ 48±11 (0.31±0.08 rem. act.) |
| **23** | 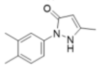 | 0.38±0.02 | inactive (30 µM) toxic (≥ 50 µM) |
| **24** | 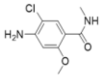 | 0.44±0.03 | $IC_{50}$ 36±9 (0.31±0.08 rem. act.) |

B

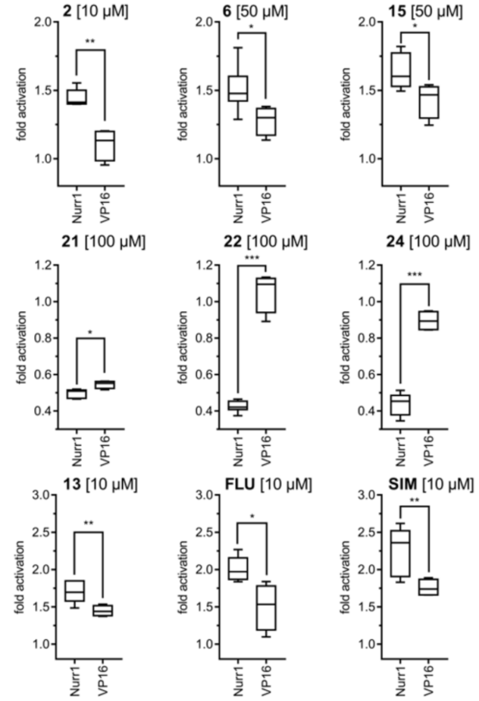


C

| drug INN (related fragement) | structure | Nurr1 modulation |
|---|---|---|
| **Astemizole** (from **2**) | 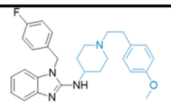 | inactive (10 µM) toxic (≥ 30 µM) |
| **Mizolastine** (from **2**) | 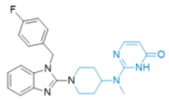 | inactive (30 µM) toxic (≥ 100 µM) |
| **Chlorpyramine** (from **6**) | 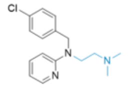 | inactive (100 µM) |
| **Mirtazapine** (from **6**) | 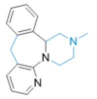 | inactive (100 µM) |
| **Procaterol** (from **15**) | 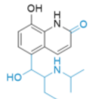 | inactive (100 µM) |
| **Chloroxine** (from **15**) | 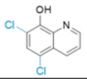 | toxic (≥ 10 µM) |
| **Sulfadoxine** (from **21**) | 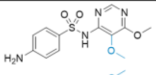 | $IC_{50}$ 47±18 µM (0.57±0.10 rem. act.) |
| **Sulfodimethoxine** (from **21**) | 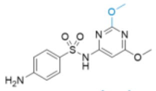 | $IC_{50}$ 14±6 µM (0.38±0.11 rem. act.) |
| **Dolasetron** (from **22**) | 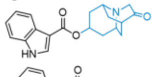 | inactive (100 µM) |
| **Ondansetron** (from **22**) | 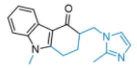 | inactive (100 µM) |
| **Cisapride** (from **24**) | 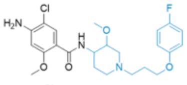 | inactive (100 µM) |
| **Metoclopramide** (from **24**) | 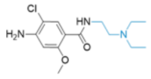 | inactive (100 µM) |

**Supplementary Figure 2**. Follow up of the primary drug fragment screen for Nurr1 modulation. (A) 24 primary screening hits further considered and their Nurr1 modulatory activity in a Gal4-Nurr1 hybrid reporter gene assay. Reporter activity in primary screen is mean±SD reporter activity, n=2. Nurr1 modulation: only activities validated against Gal4-VP16 are reported. $EC_{50}$ or $IC_{50}$ values are mean±S.E.M.; n≥3. Maximum activation or remaining activity refers to the maximum reporter activation or repression efficacy compared to DMSO (0.1%) treated cells. Toxic false positive hits from the initial screen were not further investigated. (B) Control experiments employing a Gal4-VP16 hybrid receptor were performed to confirm or refute Gal4-Nurr1 mediated activity in the cellular hybrid reporter gene assay. Boxplots show: center line, median; box limits, upper and lower quartiles; whiskers, min/max; $n \geq 4$. * $p < 0.05$, ** $p < 0.01$ *** $p < 0.001$ (t-test). (C) Nurr1 modulatory activity of fragment derived drugs on Nurr1 in a Gal4-Nurr1 hybrid reporter gene assay. Structural extensions compared to the underlying fragments are shown in blue. Only statins and the antibiotics sulfadoxine and sulfadimethoxine retained the Nurr1 modulatory activity of their fragment precursors **13** and **21** (see also Figure 1B). $IC_{50}$ values are mean±S.E.M.; n≥3.

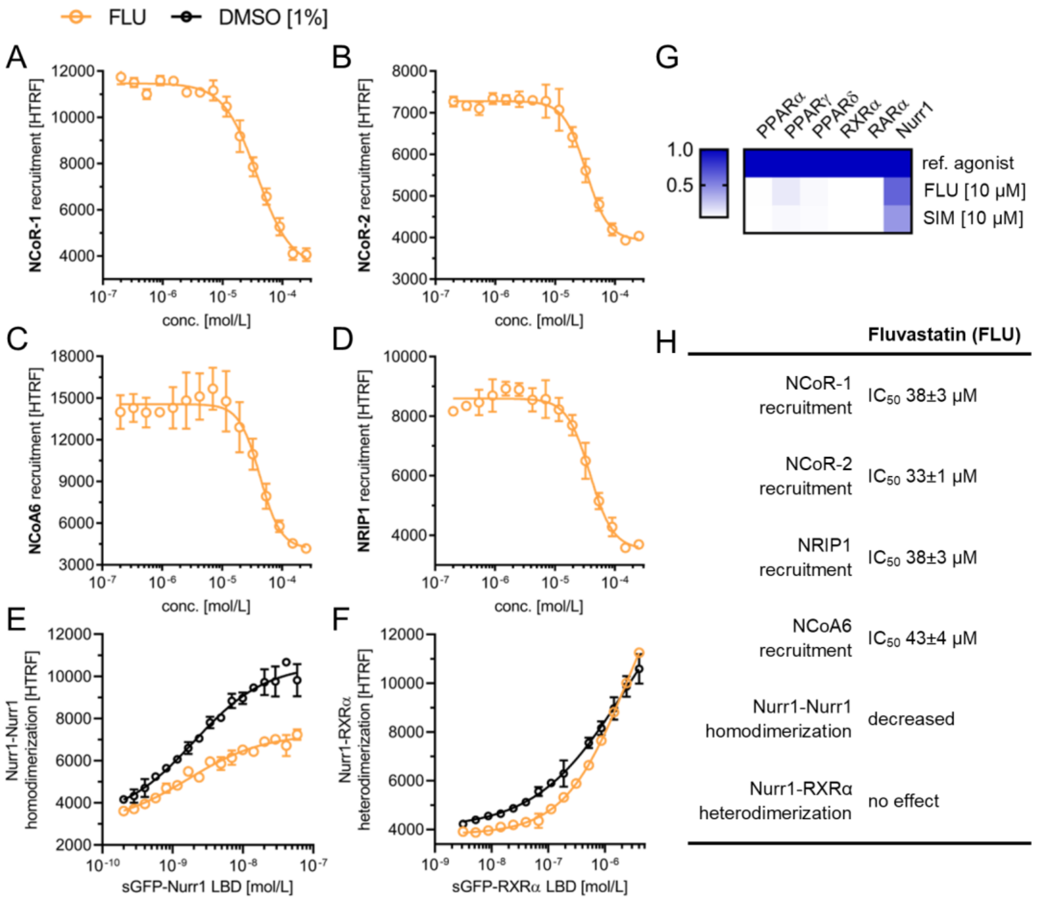


**Supplementary Figure 3**. Effects of fluvastatin (FLU) on co-regulator interactions and dimerization of Nurr1. (A-D) Fluvastatin displaced NCoR-1 (A), NCoR-2 (B), NCoA6 (C) and NRIP1 (D) from the Nurr1 LBD. (E, F) Fluvastatin decreased homodimerization of Nurr1 (E) without affecting Nurr1-RXRα heterodimerization (F). Data are the mean±SD; N=3. (G) Selectivity profiles of FLU and SIM at 10 μM on related lipid-activated transcription factors in Gal4 hbrid reporter gene assays. Heatmap shows mean relative activation compared to reference agonists at 1 μM for PPARs (α: GW7647; γ: rosiglitazone; δ: L165,041), RXRα (bexarotene), RARα (tretinoin) and 100 μM for Nurr1 (amodiaquine); n ≥ 2. (H) Summarized cell-free Nurr1 modulatory activities of FLU.

### Supplementary Methods

**Computational Methods.** *General*: Calculations were conducted in KNIME (version 3.7.2, KNIME AG, Zurich, Switzerland) and Molecular Operating Environment (MOE, version 2018.0101, Chemical Computing Group Inc. Montreal, QC, Canada) using default settings for each tool/function unless stated otherwise. Amber10:EHT was used as default force field for all calculations. *Library processing*: Analysis of the drug fragment library from Prestwick (Prestwick Chemical, Illkirch, France) was performed in KNIME using the provided SMILES strings compared to the DrugBank database (all drug structures in SDF Format, version 5.1.1, released on 2018-07-03). The RDKit extension nodes (version 4.0.1.v202002121354) were used to filter for PAINs structures and calculate features (MW, clogP, number of H-bond donors/acceptors and rotatable bonds, aromatic rings, TPSA). For hits from the primary screen, a search for parent and related drugs was performed via molecule substructure and murcko scaffold (both RDKit nodes) compared to the DrugBank database (version 5.1.1). Graph based frameworks were extracted with the MOE KNIME extension node murcko frameworks ignoring small terminal rings of size 3 or 4. Rings of size 5 to 7 atoms as well as annealed rings, bicyclo and spiro compounds of equal ring count were assigned to the same groups. Geometry in terms of linker attachment points and connectivity was ignored, only the linker length was considered. *Multiple alignment*: Molecular structures of amodiaquine, fluvastatin and pitavastatin were prepared using MOE Wash tool: protonation state dominant at pH 7; coordinates rebuild 3D; preserved existing chirality. Multiple alignment of these three compounds was performed using default settings from MOE flexible alignment tool.
